# Differences in white matter detected by *ex vivo* 9.4T MRI are associated with axonal changes in the R6/1 model of Huntington’s Disease

**DOI:** 10.1101/2023.10.02.560424

**Authors:** C. Casella, B. Kelly, A. Murillo Bartolome, B. Mills-Smith, G.D. Parker, C. Von Ruhland, Y.A. Syed, V. Dion, A.E. Rosser, C. Metzler-Baddeley, D.K. Jones, M.J. Lelos

**Affiliations:** Cardiff University Brain Research Imaging Centre (CUBRIC), School of Psychology, Cardiff University, Cardiff, UK; School of Biosciences, Cardiff University, Cardiff CF10 3AX, U.K; UK Dementia Research Institute at Cardiff University, Cardiff CF24 4HQ, UK; B.R.A.I.N unit, Neurosciences and Mental Health Institute, School of Medicine, University Hospital of Wales, Heath Park, Cardiff, CF24 4HQ, U.K

**Author notes:** Department of Perinatal Imaging and Health, School of Biomedical Engineering & Imaging Sciences, King’s College London, St Thomas’ Hospital, London, UK. **Corresponding Author**, Dr. Chiara Casella Department of Perinatal Imaging and Health, School of Biomedical Engineering & Imaging Sciences, King’s College London, St Thomas’ Hospital, London, UK.

**Keywords:** Diffusion MRI, Quantitative magnetization transfer, Huntington’s disease, White matter microstructure, R6/1 mice, Ultra-high field MRI

## Abstract

White matter (WM) volume loss has been reported in people with Huntington’s disease (HD), but the cellular basis of this deficit remains to be elucidated. To address this, we assessed ex vivo WM microstructure in the transgenic R6/1 mouse model of HD with magnetic resonance imaging (MRI) and studied the neurobiological basis of the MRI brain signals with histological and electron microscopy analyses in a separate cohort of age- and sex-matched mice. Differences in the macromolecular proton fraction (MPF) from quantitative magnetization transfer (qMT) as a proxy myelin measure, and the intra-axonal signal fraction (FR) from the composite hindered and restricted model of diffusion (CHARMED) as a proxy marker of axon density, were assessed alongside diffusion tensor imaging (DTI) parameters. A tractometry approach was employed to inspect region-specific differences across the corpus callosum (CC). Furthermore, voxel-based morphometry (VBM) and tract-based spatial statistics (TBSS) were used to explore brain-wise WM macro- and microstructure abnormalities. To gain insight into disease-associated impairments in attentional and visuospatial processing, a third cohort of age-matched mice was assessed with the 5-choice serial reaction time task (5-CSRTT). We report cognitive impairments in R6/1 mice and, by evaluating MRI and light and electron microscopy results, we show that this HD mouse model presents disruptions in axonal morphology (i.e. less complex, thinner axons) and organization (i.e. more densely packed axons). Furthermore, we show that, at least early in disease progression, R6/1 mice present a reduction in the expression or content of myelin-associated proteins without significant alterations in the structure of myelin sheaths. Finally, our findings indicate that neuroinflammation-driven glial and axonal swelling might also affect this mouse model early in disease progression. Crucially, we demonstrate the potential of FR, an in vivo estimate of axon density, as a novel MRI biomarker of HD-associated changes in WM microstructure.

## 1. Introduction

Huntington disease (HD), a neurodegenerative disorder leading to devastating cognitive, psychiatric and motor symptoms, cannot currently be cured. It is therefore a research priority to enhance understanding of how it develops and identify biomarkers to support therapy development. Recent investigations in HD have recognized alterations in the brain’s white matter (WM) as relevant pathophysiological feature of HD ^1–12^.

Specifically, WM atrophy has been shown both in animal models and in human HD carriers by histopathological post-mortem studies ^13–17^, and MRI studies ^2,4,5,11,17–21^. For example, structural neuroimaging studies of HD carriers have shown that WM atrophy can be found across several WM areas, including the corpus callosum (CC), the anterior commissure, internal and external capsules, and the cingulum. Furthermore, they suggest that WM changes happen very early in the disease course ^7,20,25–27^. The severity of WM atrophy has been shown to correlate with predicted time to symptom onset in pre-manifest patients ^7,9,22^, measures of motor dysfunction ^23^ and cognitive deficits ^19,24^.

However, the cause of WM degeneration and its involvement in the development and advancement of HD are still not well understood. Notably, WM is composed of axons as well as myelin-producing oligodendrocytes, and it is unclear whether it is the loss of axons, myelin, or both, that drives the WM volumetric loss ^8^. Accordingly, while most neuroimaging studies on WM alterations in HD have relied on measures of WM volume, WM volume loss as quantified using structural MRI is a rather unspecific marker of disease stage and progression as it is not sensitive to microstructure changes. As such, reductions in WM volume observed in structural neuroimaging studies can be the consequence of several factors, including a decrease in the number of axons because of Wallerian degeneration, a decrease in axon myelination, or a combination of both.

Although diffusion tensor MRI (DT-MRI) ^28^ has allowed a deeper understanding of WM organization at the microstructural level, DT-MRI metrics do not map specifically onto biological subcomponents of WM microstructure ^29^. It is therefore very hard to interpret changes in DT-MRI metrics in terms of specific microstructural properties. Very different configurations of, for example, axonal packing, axonal size, and myelination may generate very similar outcome measures (See Figure 2 in ^30^). Accordingly, while some DT-MRI studies have suggested that WM changes in HD are a consequence of axonal injury rather than demyelinating mechanisms ^31^, several others have indicated a role for myelin disturbance in HD pathology ^2,11,26,32–34^.

**Figure 1.**
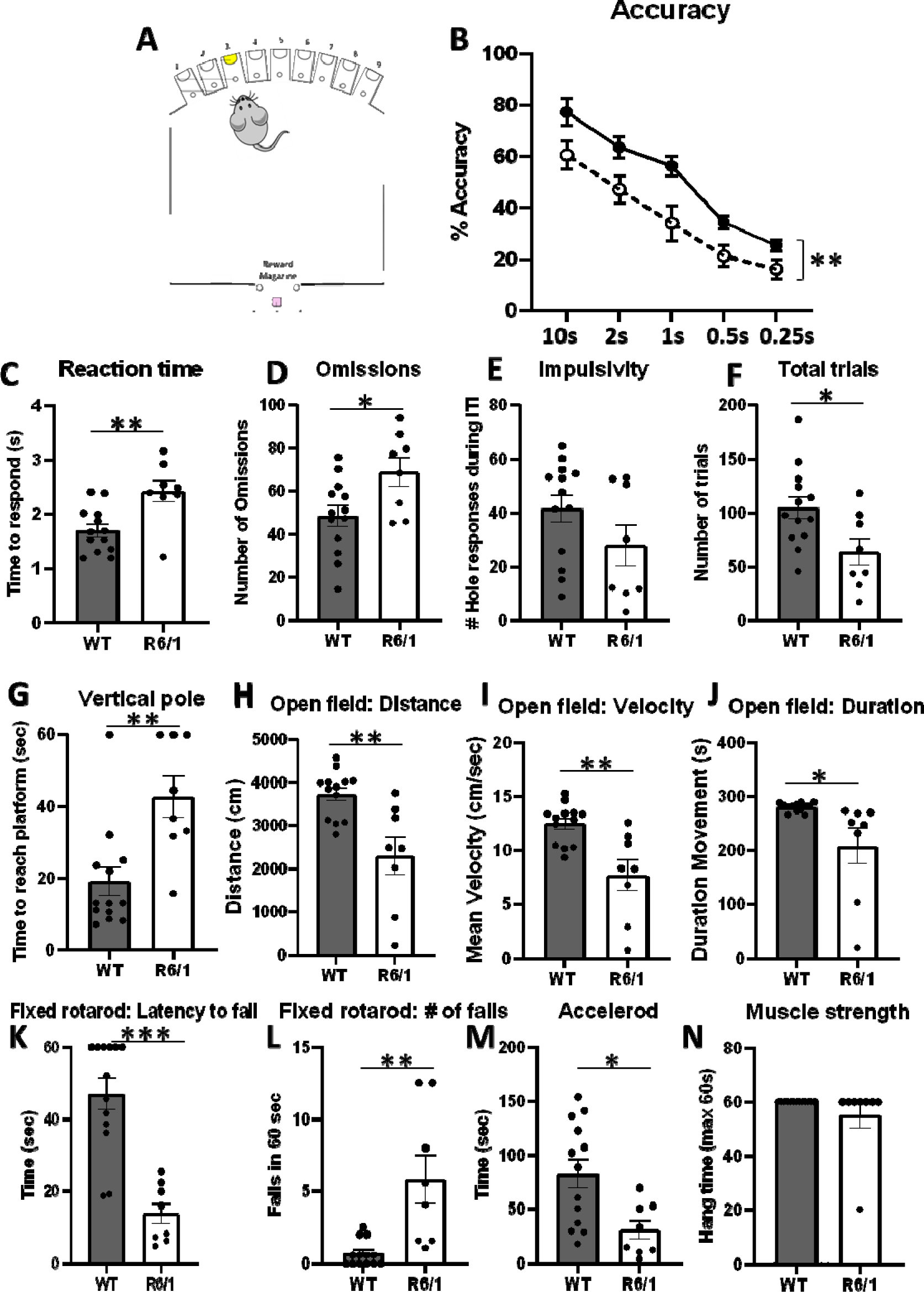
Analysis of cognitive and motor function in WT and R6/1 mice. (A) Schematic showing the 9-hole operant box apparatus with every other nosepoke hole available. (B) Impaired response accuracy in R6/1 mice is evident on the 5-CSRTT, even when the response stimulus duration was long during training (e.g., 10s). On the 5-CSRTT, R6/1 mice reacted to the stimulus more slowly (C) and omitted more responses (D). The R6/1 mice did not display impulsivity, as assessed by the number of hole pokes during the inter-trial interval (E), but they did complete fewer trials overall (F). R6/1 mice were slower on the vertical pole to reach the platform (G). In the open field, R6/1 mice travelled less far (H), were slower (I) and spent less time moving (J). On the fixed rotarod, R6/1 mice fell off more quickly (K) and fell more times across the 60s session (L). On the accelerod, R6/1 mice fell more quickly (N), but they did not show a difference in muscular strength on the hang wire test (N).

**Figure 2.**
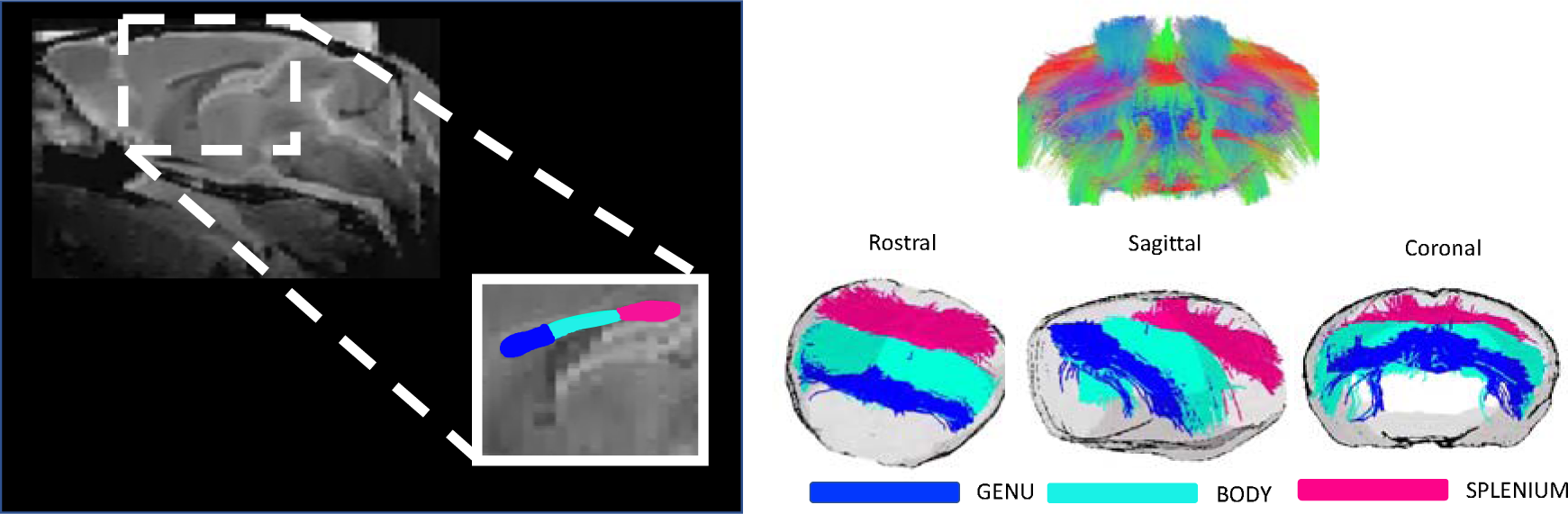
Tractography of the corpus callosum (CC) Left: Representative figure of manually delineated segments of the CC (genu, body, and splenium). Right: Coronal view of whole-brain tractography (top) and fibers travelling through the manually delineated region of interests (ROIs), in rostral, sagittal and coronal views (bottom).

In this work, we moved beyond volumetric and DT-MRI methods to obtain a greater insight into the neurobiological mechanisms underlying WM changes reported by MRI studies of HD carriers. For this purpose, we carried out a comprehensive WM microstructural assessment of the R6/1 mouse model of HD ^35,36^ using 9.4T MRI. We complemented gross measurements of WM atrophy performed with voxel-based morphometry (VBM) and fractional anisotropy (FA), axial diffusivity (AD) and radial diffusivity (RD) from DT-MRI ^37^, with measures providing increased sensitivity to tissue microstructure and biochemical composition. Specifically, we measured the macromolecular proton fraction (MPF) from quantitative magnetization transfer (qMT) ^38^ as a proxy measure of myelin, and the restricted diffusion signal fraction (FR) from the composite hindered and restricted model of diffusion (CHARMED) ^39^ as a proxy measure of axon density ^29^, in an attempt to disentangle the contribution of changes in axon microstructure versus myelin to HD pathology.

The R6/1 mouse model of HD expresses exon 1 of the human HD gene, with around 115 CAG repeats ^35^. This line was amongst the first to be developed ^35^, and presents a progressive phenotype and neuropathology, which occur early in life, with a reduced lifespan. Specifically, R6/1 mice exhibit age-related progressive changes in both motor and cognitive performance from as early as 2 months of age ^40^ and age-related neuronal loss, brain atrophy and mHTT accumulation ^41^.

Despite evidence for grey matter (GM) pathology in R6/1 mice ^42,43^, not many studies have performed an MRI characterization of WM alterations in this model. One previous study looking at gross volumetric changes in R6/1 mice, reported sparing of WM in this model, with no detectable differences in CC volume at either 9 or 17 weeks of age ^43^. On the other hand, a recent *ex vivo* MRI study by Gatto and colleagues ^42^ reported alterations in WM microstructure metrics in the callosal genu of 11 weeks old R6/1 mice. These findings suggest that, although gross WM volumetric changes are not yet detectable in these mice early in HD progression, WM microstructure is impaired. However, their histological analysis included markers of inflammation in GM, leaving cellular changes in WM alteration unexplored.

In the present study, the assessment of both macrostructural and microstructural changes in WM was performed with ex vivo MRI on the brains of 4 month old mice (N=7 WT and N=8 R6/1), with the aim of increasing understanding of the intricate cellular changes happening early in disease progression. Post-mortem imaging enabled longer scan times and limited the potential challenges commonly affecting live rodent scanning, such as movement artefacts. This in turn allowed data to be acquired with higher resolution, signal-to-noise and contrast-to-noise ratio ^44^.

We used a separate cohort (N=13 WT and N=8 R6/1) to characterise disease-associated functional impairments in this model at 4 months of age. Specifically, we assessed attentional and visuospatial processing using the 5-choice serial reaction time task (5-CSRTT), as well as a range of simple motor tasks.

Finally, to gain insight into the neurobiological basis of imaging results, we used separate age- and sex-matched cohorts of mice to carry out light microscopy analysis of axons and myelin across all segments of the CC (N=9 WT and N=9 R6/1) and transmission electron microscopy (N=3 and N=3) analyses, to more precisely measure changes in axonal size, axonal density and myelin thickness. By leveraging the increased biological specificity obtained with this approach, we hoped to better inform previous knowledge and future research in the human condition.

## 2. Materials and Methods

### 2.1. Mice

All experimental procedures in this study (Table 1) followed protocols in accordance with the United Kingdom Animals (Scientific Procedures) Act of 1986. All experimental procedures performed on mice were approved by Cardiff University Ethical Review Process Committee and carried out under Home Office License P49E8C976.

**Table 1.** Summary of assessments carried out in this study and sample description.

| Sample description | Assessment |
| --- | --- |
| 7 WT and 8 R6/1 female mice<br>tested at 4 months of age | <u>Ex vivo MRI:</u><br>▪ Tractometry of the CC (FA, AD, RD, FR and MPF)<br>▪ TBSS (FA, AD, RD, FR and MPF)<br>▪ VBM (anatomical data) |
| 13 WT and 8 R6/1 mixed-sex mice tested at<br>1-4 months of age | <u>Assessment of cognitive function:</u><br>▪ 5-CSRTT<br><u>Assessment of motor function:</u><br>▪ Balance beam<br>▪ Fixed rotarod<br>▪ Accelerated rotarod<br>▪ Vertical pole<br>▪ Locomotor activity<br>▪ Muscular strength |
| 9 WT and 9 R6/1 female mice tested at 4<br>months of age | <u>Light microscopy:</u><br>▪ NFL and MBP CC thickness and area fraction |
| 3 WT and 3 R6/1 female mice tested at 4<br>months of age | <u>Electron microscopy:</u><br>▪ Axon diameter of myelinated axons<br>▪ G-ratio of myelinated axons |

Twelve (12 weeks old) female hemizygous R6/1 mice and wildtype littermates were purchased from Jackson Laboratories (Jax^®^, Bar Harbour, Maine, U.S.A.) to be scanned at 16 weeks of age. A final sample of 7 WT and 8 R6/1 mice was examined in this study, due to fixation artifacts present in the other samples, which made them inadequate for quantitative data analysis ^45^.

An age- and sex-matched cohort of R6/1 and WT littermate mice was used for electron microscopy (N=3 WT and N=3 R6/1). For light microscopy, an age- and sex-matched cohort that consisted of N=9 WT and N=9 R6/1 were used, but sections from a 1:12 series were only included for analysis if they fell within the boundaries determined for the genu, body and splenium (see Section 2.5).

For motor and cognitive testing, an age- and mixed-sex cohort of mice consisting of WT (N=13) and R6/1 (N=8) was used. All these mice were injected with a control viral vector at P2, but subsequent to the behavioural testing, it was found that the vector lacked efficacy.

Prior to the experiment, all animals were housed in age- and sex-matched groups of between 1 and 5 mice, with mixed genotypes. Mice were subject to a 12-hour light:12-hour dark cycle with controlled room temperature (21 ± 3 °C) and relative humidity (60 ± 3%). Each cage contained modest environmental enrichment including play tunnels and nesting material. All animals were weighed on a weekly basis to monitor general health.

### 2.2 Assessment of visuospatial and attention function in the 5-CSRTT, and simple motor function

Mice were trained on the 5-choice serial reaction times task (5-CSRTT), starting at 1 month of age. They were food restricted 5 days before training commenced and the 5-CSRTT testing concluded when mice were ∼3 months old. The 5-CSRTT task requires the mouse to attend to the array until a stimulus light is briefly presented. Successful nose poke in the operandum where the stimulus flash occurred results in milkshake reward. Failure to poke in the correct hole results in a brief timeout, during which time the house light is presented briefly, before another trial commences. This task measures visuospatial attention during the interval prior to onset of the stimulus, when the mouse is required to attend to the 5-hole array to detect the flash of the stimulus light. During this same period, impulsive behaviours can be detected if mice respond inappropriately, rather than waiting for the stimulus presentation. Accuracy refers to the number of correct responses made out of the total number of trials that commenced. The reaction time is the duration of time between the stimulus onset and the mouse successfully responding in the nose poke operandum. Omissions refer to trials in which the stimulus light flashes, but no response is made during the 10 second limited hold period. Impulsivity refers to the number of nose pokes that occur during the interval prior to stimulus onset. Total trials refers to the total number of trials successfully completed during the test session.

Training was conducted as described in ^46^. See Figure 1A. In brief, mice were initially trained to retrieve strawberry milkshake reward from the magazine, followed by training to respond in the central nosepoke of the array. During the initial training stages, the stimulus light was maintained for 10s to support learning, and this was gradually reduced to 2s, 1s, 0.5s and finally 0.25s stimulus duration.

From 3-4 months old, mice were maintained on an ad lib diet and tested on a range of simple motor tests, as described in ^40^. Four trials of balance beam were conducted and data were averaged across the trials. Two trials of fixed rotarod (12 rotations per minute) were conducted and the data were analysed across these two trials. Data consisted of the mean latency to fall and the number of falls in a 60s period. Two trials of the accelerated rotarod were run and data were averaged across these two trials. Two trials of vertical pole were conducted and data were averaged across these two trials. Locomotor activity was also tracked in the open field using Ethovison over 5 minutes. Finally, muscular strength was assessed using the wire hang test for a maximum of 60s.

#### Statistical analysis

All data were analysed using GraphPad Prism 9. The accuracy data from 5-CSRTT data wereanalysed by AONVA with stimulus duration (10s-0.25s) as a within subjects factor andgenotype as a between subjects factor. The remaining measures (reaction time, impulsivity,omissions and total trials) were analysed by independent subjects t-test. The vertical poletest data were analysed by independent subjects t-test. For the open field test the threemeasures (distance, velocity and duration) were analysed by independent subjects t-test.For the fixed rotaroad, the two measures (Time (latency to fall) and number of falls in 60s)were analysed by independent subjects t-test. The accerod data of time (latency to fall) wasanalysed by independent subjects t-test. For the grip test of muscle strength, the data wereanlysed by independent subjects t-test.

### 2.3. Perfusion

For imaging and light microscopy, mice were terminally anaesthetised *via* intraperitoneal injection of 0.3 ml Euthatal and then perfused through the left ventricle with approximately 60 ml of phosphate buffered saline (PBS). This was followed by infusion with about 150ml of 4% formaldehyde in PBS (pH 7.3) at a flow rate of 30ml/min. The temperature of the perfusates was maintained at room temperature and the pH of the formaldehyde was 7.2–7.4. After decapitation, the skulls were de-fleshed and post-fixed in 4% formaldehyde in PBS overnight. They were then transferred to a 25% sucrose solution and stored at 4°. For electron microscopy, a similar procedure was followed, except that the perfusate contained 2.5% Glutaraldehyde + 2.5% paraformaldehyde in PBS.

### 2.4. Ex vivo MRI Assessment of R6/1 brains

#### Tissue preparation

Brains were scanned in-skull, in order to preserve neural structures. The skulls were soaked in PBS and washed daily for three days, to regain some signal due to tissue rehydration ^47^. The skulls were then carefully wiped with tissue paper, and immersed in Galden, a proton-free susceptibility-matching fluid in a 15ml syringe. The use of a syringe allowed any residual air bubbles to be pushed out, which might otherwise have affected MRI measurements. Immediately after scanning, skulls were returned to PBS and washed for three days, before being stored in a 25% sucrose −0.1% sodium azide solution at 4°, to ensure tissue preservation.

#### Data acquisition

MRI acquisition was conducted *ex vivo* on a 9.4 Tesla (20 cm) horizontal-bore animal system (Bruker Biospin, Germany). This was equipped with BGA12-S (12cm inner bore size, integrated shims) gradients. A transmit 1H 500 Watt echo-planar imaging (EPI) volume coil was used with a phased array 4-channel surface coil and Paravision software (version 6.1, Bruker Biospin) were used for data acquisition.

The magnetic field homogeneity was optimized with a localized shimming procedure (Fastmap, Bruker Biospin) on a volume of interest placed in the centre of the brain. A PRESS-waterline sequence (Bruker BioSpin) was used with outer volume suppression without water suppression to evaluate water line width (TR/TE=2500/20ms respectively) to assess the shim performance and the peak line width of the water signal. Iterations were repeated until all line widths < 40 Hz.

The acquisition protocol consisted of a T_1_-weighted FLASH sequence, a multi-shell dMRI acquisition for DTI and CHARMED ^28,39^, and an MT-weighted (MT-w) T_1_ FLASH sequence. Additionally, the longitudinal relaxation rate of the system was estimated by acquiring T_1_-maps using T_1_-weighted FLASH images. A description of the acquisition parameters for each sequence is provided in Table 2.

**Table 2.**
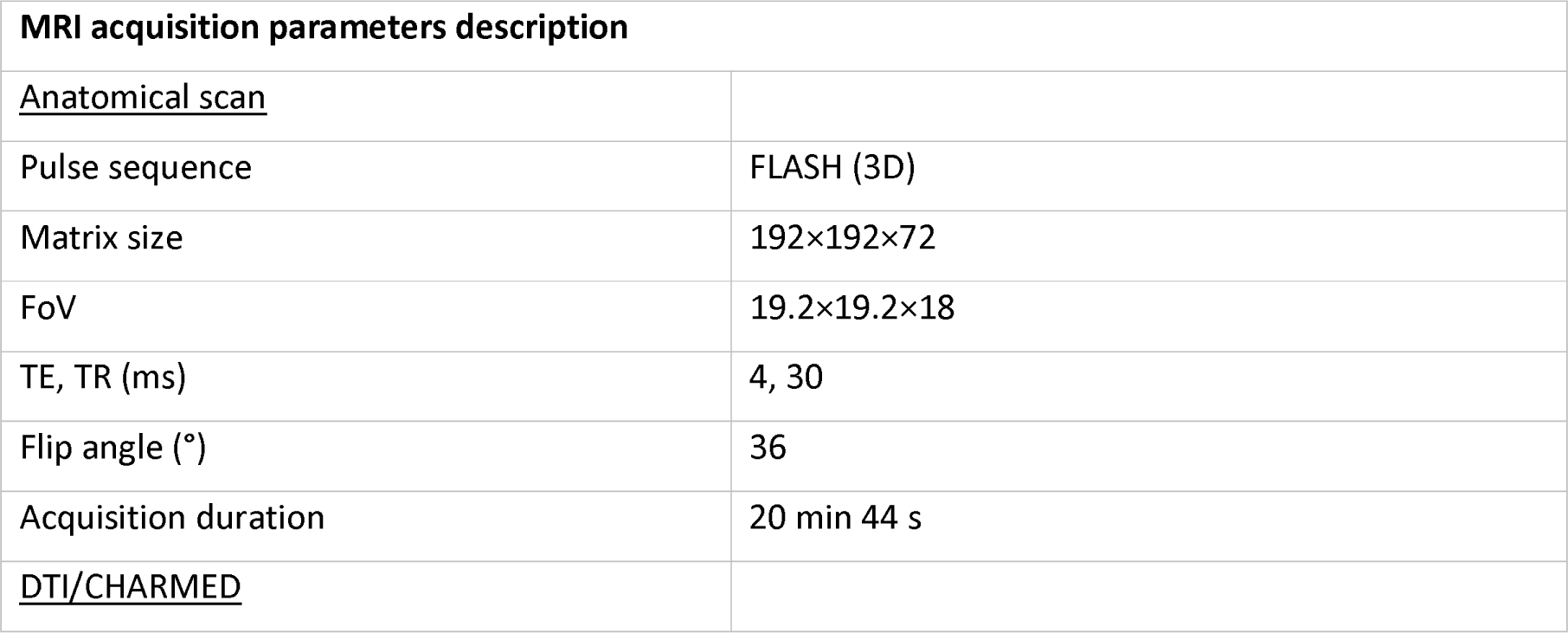
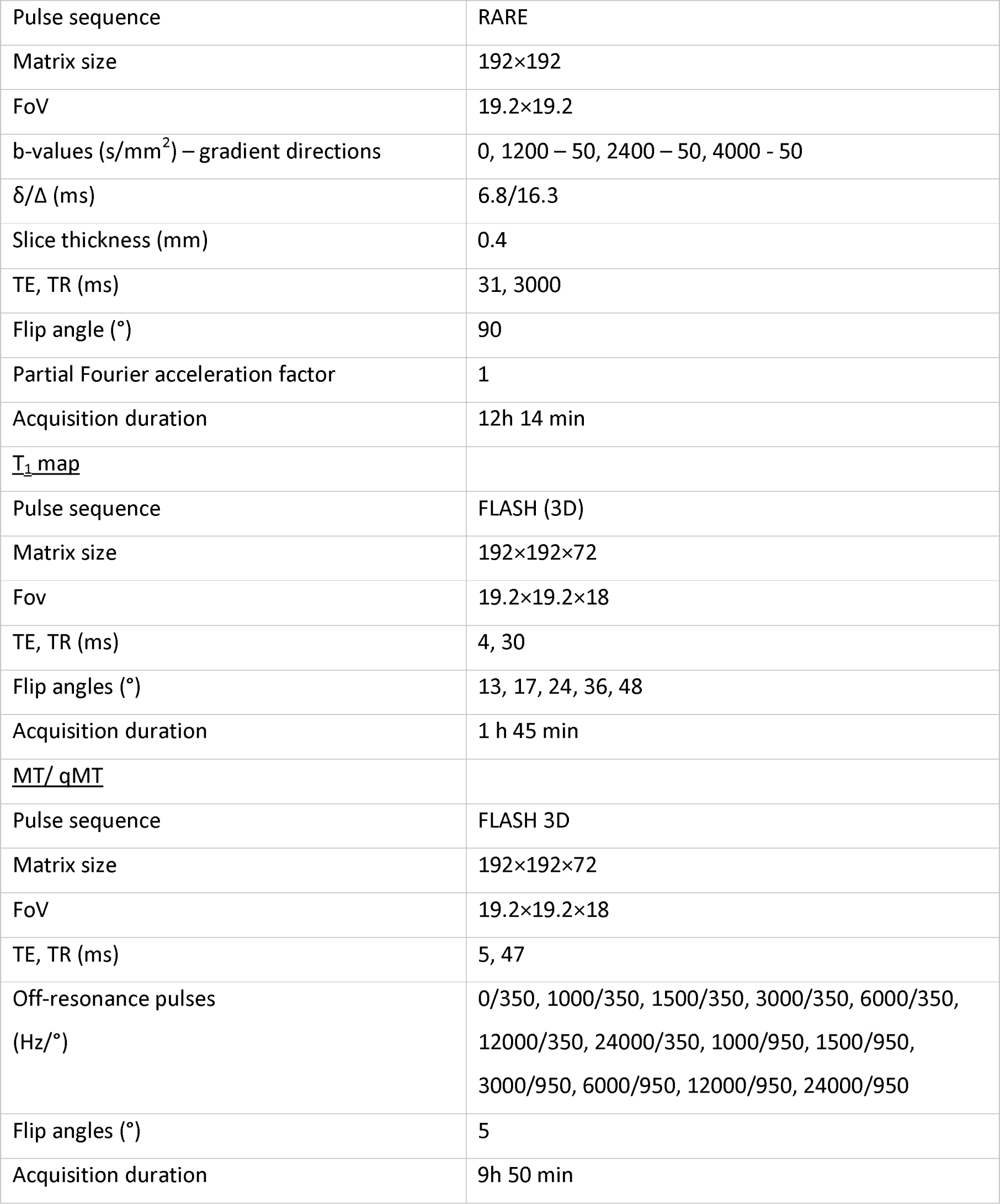
Scan parameters. All sequences were acquired at 9.4 Tesla. For each of the sequences, the main acquisition parameters are provided. MT: magnetization transfer; qMT: quantitative magnetization transfer. FoV: field of view; TE: echo time; TR: repetition time.

| MRI acquisition parameters description |  |
| --- | --- |
| <u>Anatomical scan</u> |  |
| Pulse sequence | FLASH (3D) |
| Matrix size | 192×192×72 |
| FoV | 19.2×19.2×18 |
| TE, TR (ms) | 4, 30 |
| Flip angle (°) | 36 |
| Acquisition duration | 20 min 44 s |
| <u>DTI/CHARMED</u> |  |
| Pulse sequence | RARE |
| Matrix size | 192×192 |
| FoV | 19.2×19.2 |
| b-values (s/mm<sup>2</sup>) – gradient directions | 0, 1200 – 50, 2400 – 50, 4000 – 50 |
| $\delta/\Delta$ (ms) | 6.8/16.3 |
| Slice thickness (mm) | 0.4 |
| TE, TR (ms) | 31, 3000 |
| Flip angle (°) | 90 |
| Partial Fourier acceleration factor | 1 |
| Acquisition duration | 12h 14 min |
| <u>T<sub>1</sub> map</u> |  |
| Pulse sequence | FLASH (3D) |
| Matrix size | 192×192×72 |
| Fov | 19.2×19.2×18 |
| TE, TR (ms) | 4, 30 |
| Flip angles (°) | 13, 17, 24, 36, 48 |
| Acquisition duration | 1 h 45 min |
| <u>MT/ qMT</u> |  |
| Pulse sequence | FLASH 3D |
| Matrix size | 192×192×72 |
| FoV | 19.2×19.2×18 |
| TE, TR (ms) | 5, 47 |
| Off-resonance pulses<br>(Hz/°) | 0/350, 1000/350, 1500/350, 3000/350, 6000/350,<br>12000/350, 24000/350, 1000/950, 1500/950,<br>3000/950, 6000/950, 12000/950, 24000/950 |
| Flip angles (°) | 5 |
| Acquisition duration | 9h 50 min |

#### Image processing

Skull-stripping was performed using the Rodent Bet Brain Extraction Tool, a modified version of the Brain Extraction Tool (BET; FSL v5.0) that can process rodent brains ^48^.

##### Anatomical data

Processing of T_1_ anatomical images was performed using SPM8 (Wellcome Trust Institute of Neurology, University College London, UK, www.fil.ion.ucl.ac.uk/spm) with the SPMMouse toolbox (http://spmmouse.org) for animal brain morphometry ^49^. This toolbox extends SPM’s functionality with affine registration priors for mouse brains.

Specifically, a previously described mouse brain atlas ^50^ was used to register images from the brains of R6/1 mice with those of WT littermate controls. Following approximate manual registration using the SPM interface, images were bias-corrected and the affine priors supplied were used to register the images to the tissue probability maps. The registered images were then used to obtain GM, WM and CSF segmentations. The resulting WM segmentations were output in rigid template space and DARTEL ^51^ was used to create both non-linearly registered maps for each subject, and common templates for the cohort of animals. The warped WM portions for each mouse brain were modulated with the Jacobian determinant from the DARTEL registration fields to preserve tissue amounts, and smoothed with a Gaussian kernel of 400μm to produce maps for analysis ^52^.

##### Diffusion data

Pre-processing of DWI data was performed with various tools including FSL ^53^, MRtrix3 ^54^, ExploreDTI (v.4.8.3) ^55^, and ANTS ^56^. Each diffusion dataset was denoised ^57^ and corrected for field distortion ^58^ and Gibbs ringing artefacts ^59^. Diffusion tensors were estimated using non-linearly weighted least squares (for b < 1500 s/mm^2^ data) providing the following quantitative scalar measures: FA, AD and RD. Motion and distortion artefacts in the CHARMED data were corrected according to the extrapolation method described in ^60^. FR maps ^39^ were computed by fitting the CHARMED model to the DWI data, with an in-house software coded in MATLAB (The MathWorks, Natick, MA).

##### qMT data

MT-w images were corrected for Gibbs ringing artefacts ^59^ and co-registered to the MT-w volume with the most contrast using an affine registration (FLIRT, 12 degrees of freedom, mutual information). Subsequently, the MT-w images and T_1_-maps were modelled by the two-pool Ramani’s pulsed MT approximation ^61^. This provided MPF maps, which were nonlinearly registered to the b=0s/mm^2^ image using the MT-w volume with the most contrast as a reference, using FNIRT ^62^. Accuracy of all registrations was confirmed by visual inspection.

#### Tractography of the CC

Multi-shell, multi-tissue, constrained spherical deconvolution (MSMT-CSD) ^63^ was applied to the pre-processed images to obtain voxel-wise estimates of the fiber orientation density functions (fODFs) ^64–66^ with maximal spherical harmonics order lmax=8. The fODFs were generated using a set of 3-tissue group-averaged response functions ^67^. Seed points were positioned at the vertices of a 0.2×0.2×0.2mm grid superimposed over the image. The tracking algorithm interpolated local fODF estimates at each seed point and then propagated 0.05mm along orientations of each fODF lobe. This process was repeated until the minimally subtending peak magnitude fell below 0.05 or the change of direction between successive 0.05mm steps exceeded an angle of 40°. Tracking was then repeated in the opposite direction from the initial seed point. Streamlines whose lengths were outside a range of 2mm to 30mm were discarded.

To assess regionally specific effect of HD on the CC, tractography was performed in three different callosal segments (genu, body and splenium). 3D fibre reconstructions were performed interactively in the native space of each mouse in FiberNavigator ^68^ (Figure 2), using a combination of include and exclude regions of interest (ROIs), according to the following protocols:

##### Genu

Two ROIs were placed anterolateral to the most rostral portion of the corpus callosum in each hemisphere. This approach was used to capture the anteriorly arching fibres of the genu ^69^. An exclusion ROI was used to exclude streamlines extending posteriorly to the genu on the sagittal plane, which make up the body of the CC.

##### Body

Two ROIs were placed ventral to the location of the cingulum and medial to the lateral ventricles (one in each hemisphere) ^69^.Exclusion ROIs were used to exclude the genu and splenium (i.e., the anterior and posterior sections of the CC, respectively).

##### Splenium

Two ROIs were placed posterolateral to the most caudal section on the CC in the left and right hemisphere ^69^. An exclusion ROI was used to remove streamlines extending anteriorly to the splenium.

#### Statistical analysis

Statistical analyses were carried out using RStudio v1.1.456 ^70^, MATLAB (The MathWorks, Natick, MA), SPSS version 20119 ^71^, FSL ^53^ and the SPMMouse toolbox (http://spmmouse.org) for animal brain morphometry ^49^.

##### Tractometry of the CC

Microstructure differences were assessed in each of the three callosal segments. First, by taking each quantitative metric map (each registered to the b=0s/mm^2^ image during pre-processing), samples of each metric were obtained at each vertex of the reconstructed segments, and segment-specific medians were derived for FA, AD, RD, FR and MPF in MRtrix3 ^54^. Next, the overall mean was calculated, so that each dataset comprised 5 MRI-derived measures, mapped along 3 callosal segments.

##### Investigation of group differences in callosal microstructure

Two-way robust mixed ANOVAs were run for each metric (i.e., FA, RD, AD, FR, MPF), using the “bwtrim” R function from the WRS2 package ^72^. This implements robust methods for statistical estimation and therefore provides a good option to deal with data presenting small sample sizes, skewed distributions and outliers ^73^. Group was the between-subject factor and segment was the within-subject variable. Given that this robust method does not readily allow for post-hoc tests, we adopted a manual approach to further investigate significant effects. Trimmed means for each segment were calculated using a trimming percentage consistent with the one used in the primary analysis (20%). After obtaining the trimmed means, pairwise comparisons were conducted with paired-samples t-tests, and p-values corrected for multiple comparisons using the false discovery rate (FDR). Across all analyses, outliers that were ± 3 standard deviations from the mean were removed.

##### Automatic evaluation of WM atrophy using VBM

A general linear model to assess group differences in WM volume was evaluated using SPMMouse ^49^. ICV was included as a covariate of no interest as this was shown to improve estimation of volume differences in previous literature ^49^. ICV was calculated as the sum of voxels identified as GM, WM and CSF in native space for each animal, to model out the effect of different brain sizes. The sum of the WM tissue probability maps was used as explicit mask in the analysis. An adjusted p-value was calculated to control the voxel-wise FDR ^74^.

##### Assessment of brain-wise group differences in WM microstructure using TBSS

To perform a whole-brain analysis of WM microstructure changes associated with HD, the TBSS protocol ^75^, modified for use in rodents ^76^ was used. All FA maps were submitted to a free-search for a best registration target in order to minimize the image warping required for each brain volume. Specifically, each volume was first registered to every other volume, and the one requiring minimum transformation to be registered to other volumes was selected as the best registration target. This target was used as a template into which the registration was performed. Following registration, a mean FA map was calculated, thinned to represent a mean FA skeleton, and an optimal threshold of 0.2 was applied to the mean FA skeleton to create a binary WM skeleton mask. The application of such threshold allowed to exclude from further analysis areas of the brain of low FA, including peripheral small tracts, where there may be high between-subject variability and GM, and it is therefore unsafe to assume good tract correspondence across subjects ^75^.

The local FA-maximum was projected onto this WM skeleton. Subsequently, the voxel location of the local FA maximum was employed to project the respective AD, RD, FR and MPF values from that voxel onto the skeleton. Differences in microstructure measures between the two groups were assessed using voxel-wise independent t-tests (assessing areas where WT > R6/1 and R6/1 > WT). The randomize function (part of FSL) was used, together with the TFCE algorithm ^77^, generating cluster-size statistics based on 1000 random permutations. For multiple comparison correction, FDR correction was used with a threshold of p < 0.05.

### 2.5. Cytoarchitectural assessment of the CC using immunohistochemistry

#### Tissue processing

WT and R6/1 brains were frozen on a sledge-microtome (Leitz, Wetzlar), cut into 40μm coronal sections, and collected in 12 parallel series. Sections were stored in ethylene glycol-based cryoprotectant at –20C until processing. For immunostaining, 1:12 series of sections were quenched for 5 min using 10% H_2_O_2_ (VWR, West Sussex, UK) and methanol (Sigma-Aldrich, Dorset, UK). Sections were blocked in 3% serum in Triton-X and Tris-buffered saline (TxTBS) for 1 h and incubated in a solution of TxTBS, 1% serum and primary antibodies raised against either myelin basic protein (MBP; 1:1000, Santa Cruz, cat. # SC1394R) or 68kDa Neurofilament (NFL; 1:500, Abcam, cat. # ab72997). Incubation in biotinylated-secondary antibody (1:200) was conducted for 2 h, then sections were incubated using an ABC kit (Vector Laboratories Ltd, Peterborough, Cambridgeshire) for 2 h. Proteins were visualised using 3–3′-diaminobenzadine (DAB), before mounting on to double-subbed 1% gelatinised slides (Thermo Scientific, Menzel Gläser). After dehydration and delipidisation in 100% xylene, slides were coverslipped using DPX mountant (Thermo Scientific, Raymond Lamb, Leicestershire, UK).

#### Determination of CC regions of interest (ROIs)

To remain consistent with the DTI data, the CC was divided into three equal sections along its rostral-caudal axis to represent the genu, the body, and the splenium, mentioned rostral-caudal respectively. Bregma co-ordinates according to the Paxinos and Franklin Mouse Brain Atlas ^78^ were used: rostral genu 1.18mm to 0.86mm, caudal genu 0.14mm to 0.02mm, rostral body −0.1mm to −0.34mm, caudal body −0.94mm to 1.22mm and the splenium −1.3mm to −2.54mm. Three ROIs were taken from each brain hemisphere as well as 1 Medial ROI (Figure 6A). For full brain slices, ROIs were placed in identical locations on each hemisphere where their quantifications were then averaged.

#### Data collection

Bright field LM was used under identical conditions to visualise NFL and MBP immunoreactivity using the Olympus BX50 light microscope. All images of the genu, body and splenium were captured at 100x magnification with MicrosoftVIS software. The viewing field, intensity and aperture measurements remained consistent throughout imaging to ensure the data processing was indistinguishable. For image quantification, using ImageJ software (NIH, v 1.53r), all images were quantified according to two different measurements, CC thickness (μm) and area fraction (AF; i.e. the percentage of area with positive immunostaining). Images were converted to 8-bit greyscale prior to measurements being taken. For the analysis of the genu, body and splenium, all ROIs for each medial-lateral position were averaged together to determine a representative measurement (Figure 6A).

#### Statistical analysis

Statistical analysis was conducted using IBM SPSS Statistics 27. ANOVAs were conducted on data collected along the mediolateral axis, with the between-subjects factor of Genotype (WT vs R6/1) and the within-subjects factor of Region (Medial, Central-Medial, Central-Lateral and Lateral).

### 2.6. Ultra-structural analysis of axons in the CC using transmission electron microscopy

#### Tissue processing

1mm thick slices in the central-medial region of the CC body segment were post-fixed for 2 hours in 2% (w/v) aqueous osmium tetroxide, block stained for 2 hours in 2% aqueous uranium acetate, dehydrated through graded isopropanol (50%, 70%, 90% and 2 x 100%) and 3 x propylene oxide for 15 minutes each and infiltrated with TAAB embedding resin (50% in propylene oxide, 4 x neat resin for 1 hour each). Samples were placed into embedding moulds containing fresh resin and cured for 24 hours at 60C.

For LM, semithin (0.5µm) sections were collected onto droplet of distilled water on glass slides, dried on a hot plate, stained with 0.5% aqueous toluidine blue, and mounted in Gurr’s neutral mountant. Sections were examined with an Olympus BX51 research light microscope (Olympus Optical Co. (U.K.) Ltd, London, U.K) and images captured with a Zeiss Axiocam and Axiovision software (Carl Zeiss Vision GmbH, Hallbergmoos, Germany).

For electron microscopy, ultrathin (80-100nm) sections were collected onto 300 mesh copper grids, stained with Reynolds lead citrate (Reynolds, E. S. (1963)) and examined in a Hitachi HT7800 TEM (Hitachi High Tech Ltd., UK) at 100kV. Images captured with Radius software (EMSIS GmbH, Germany).

#### Image quantification

Five regions within the body of the CC were taken for quantification from randomly sampled 16.193μm^2^ electron micrographs. The axon diameter of myelinated fibres and g-ratio of myelinated axons (calculated as ‘inner diameter’/’outer diameter’, where the inner diameter was that of the axon and the outer diameter include the myelin sheath) were manually quantified using ImageJ (https://imagej.net). Specifically, the inner and outer diameters were measured at the narrowest point of each axon, assuming them to be cylindrical.

#### Statistical analysis

Statistical analysis was conducted using IBM SPSS Statistics 27. For the axonal data, an ANOVA was conducted with Genotype and Axon (myelinated, non-myelinated) as factors. For the remaining data, independent-samples t-tests were conducted with WT and R6/1 data.

## Results

### 3.1. R6/1 mice present with early cognitive and motor impairments

On the 5-CSRTT, R6/1 mice showed impaired accuracy to respond in the correct nosepoke operandum across the training (e.g. 10s) and test sessions [Figure 1B; F(1,19)=8.833, p<0.01]. R6/1 mice were slower than WT mice [Figure 1C; t(19)=3.333, p<0.01], omitted more responses [Figure 1D; t(19)=2.531, p<0.05] and completed fewer trials [Figure 1F; t(19)=2.511, p<0.05], but they did not show impulsivity, as measured by the tendency to respond in the nosepoke operandum during the intertrial period [Figure 1E; t(19)=1.538, p=n.s]. R6/1 mice showed less dexterity than WT mice on the vertical pole test [Figure 1G; t(19)=3.477, p<0.01]. On the open filed test, R6/1 mice travelled less distance, were slower and spent less time moving than WT mice [Figure 1H-J, min t(19)=2.757, p<0.05]. On the fixed rotarod, R6/1 mice fell more quickly and fell more often in the 60s session [Figure 1K-L; min t(19)=3.865, p=0.001]. R6/1 mice also showed motor coordination deficits on the accelerod [Figure 1M; t(19)=2.809, p<0.05], but no differences in muscular strength were detected using the wire hang test [Figure 1N, t(19)=1.296, p=n.s.].

### 3.2. Tractometry analysis suggests that R6/1 mice present with alterations in apparent myelin, based on MBP, and axon density, based on FR, across the CC

Results for this analysis are plotted in Figure 3.

**Figure 3.**
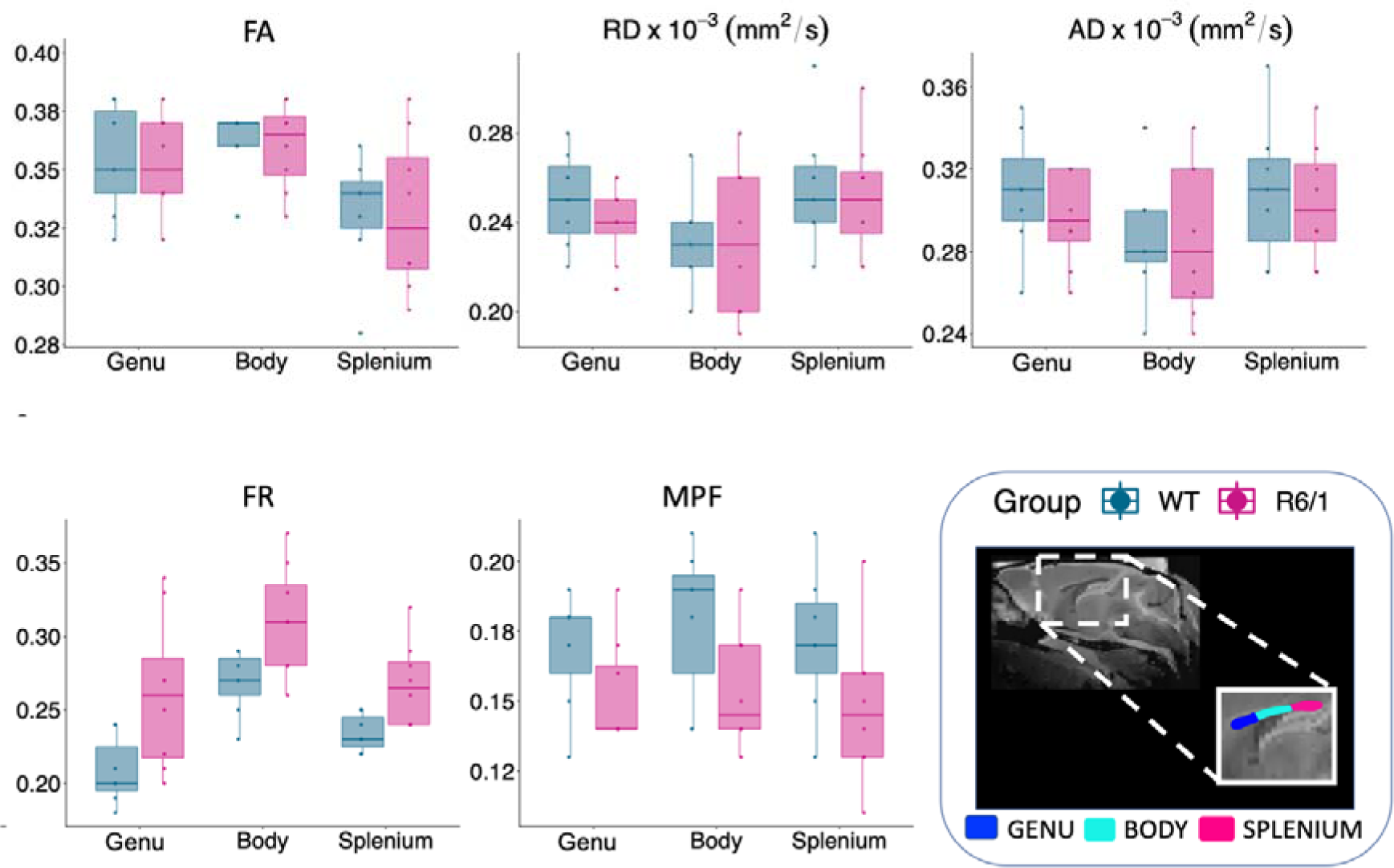
Tractometry analysis of the corpus callosum (CC). Quantification of FA, RD, AD, FR, and MPF in the genu, body and splenium of the CC. FR values were significantly higher and MPF values significantly lower in the brain of R6/1 mice across the whole CC. No significant group effects were detected for the other measures. Abbreviations: FA: fractional anisotropy; RD: radial diffusivity; AD: axial diffusivity; FR: restricted volume fraction; MPF: macromolecular proton fraction.

The mixed ANOVA for FA showed no significant effect of group [F(1, 7.4) = 0.31, p = 0.592]. On the other hand, a significant effect of segment [F(2, 6.05) = 7.51, p = 0.02] was detected, with the splenium showing significantly lower FA values compared to the body (p = 0.001). No interaction effect between group and segment was present [F(2, 6.05) = 0.009, p = 0.99].

The mixed ANOVA for RD showed no significant effect of group [Figure 3; F(1, 7.2) = 0.16, p = 0.69], but a significant effect of segment [F(2, 6.4) = 15.15, p = 0.003], with the body showing significantly lower RD values compared to the splenium (p < 0.001) and the genu (p=0.04). No interaction effect between group and segment was detected [F(2, 6.4) = 0.29, p = 0.75].

The assessment of AD values showed no significant effect of group [Figure 3; F(1, 7.79) = 0.12, p = 0.73] but a significant effect of segment [F(2, 6.2) = 10.35, p = 0.01], with the body showing significantly lower AD compared to the genu (p=0.009) and the splenium (p<0.001). The group-by-segment interaction was not significant [F(2, 6.2) = 0.44, p = 0.66].

Significant main effects of group [F(1, 5.47) = 7.21, p = 0.03] and segment [F(2, 6.36) = 43.57, p < 0.001] on FR values were detected with the mixed ANOVA. Specifically, R6/1 mice presented overall higher FR values compared to WTs across all CC regions. The CC body presented higher FR values compared to the genu (p < 0.001), the splenium presented lower FR values compared to body (p < 0.001). No significant group-by-segment interaction was detected [F(2, 6.36) = 0.24, p = 0.78].

Finally, a significant effect of group [F(1, 6.59) = 4.77, p = 0.05] was detected on MPF, with R6/1 mice showing significantly lower values. No significant effect of segment [F(2, 6.15) = 2.34, p = 0.17], nor a significant group-by-segment interaction effect [F(2, 6.15) = 0.12, p = 0.88] were detected.

### 3.3. R6/1 mice present areas of increased WM volume

Figure 4 shows regions where WM volume was significantly higher in R6/1 brains than in WT controls (p < 0.05, FDR-corrected). Increased WM volume was detected in several areas such as the posterior callosum, external capsule and olfactory bulb.

**Figure 4.**
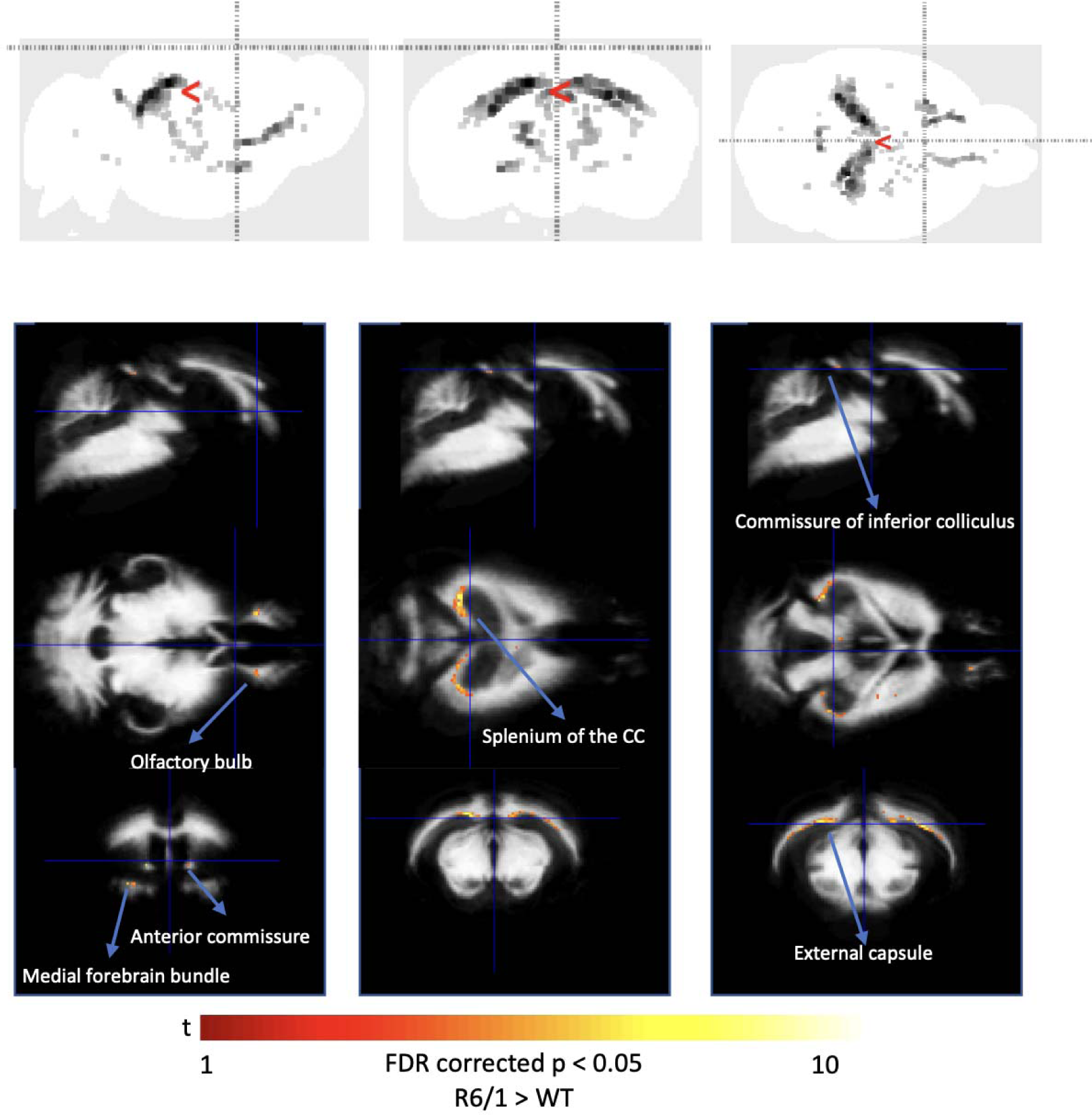
Group-differences in white matter volume. Top: Maximum intensity projections (MIPs). Bottom: Maps of the t-value (whole brain voxel analysis at p< 0.05 False discovery rate-corrected for multiple comparisons) overlayed on template white matter map, demonstrating increased volume in several areas across the brain of R6/1 mice.

### 3.4. TBSS reveals widespread increases in apparent axon density in R6/1 mice, based on FR, the restricted diffusion signal fraction

At first, a highly conservative FWE correction method was used for the TBSS analysis. This approach showed widespread increases in FR in the WM of R6/1 mice (Figure 5, top). When the voxel-wise analysis was repeated using a less conservative FDR correction of 5% (Benjamin and Hochberg, 1995), additional WM changes were detected, including more extended increases in FR and some decreases in MPF in R6/1 brains (Figure 5, bottom).

**Figure 5.**
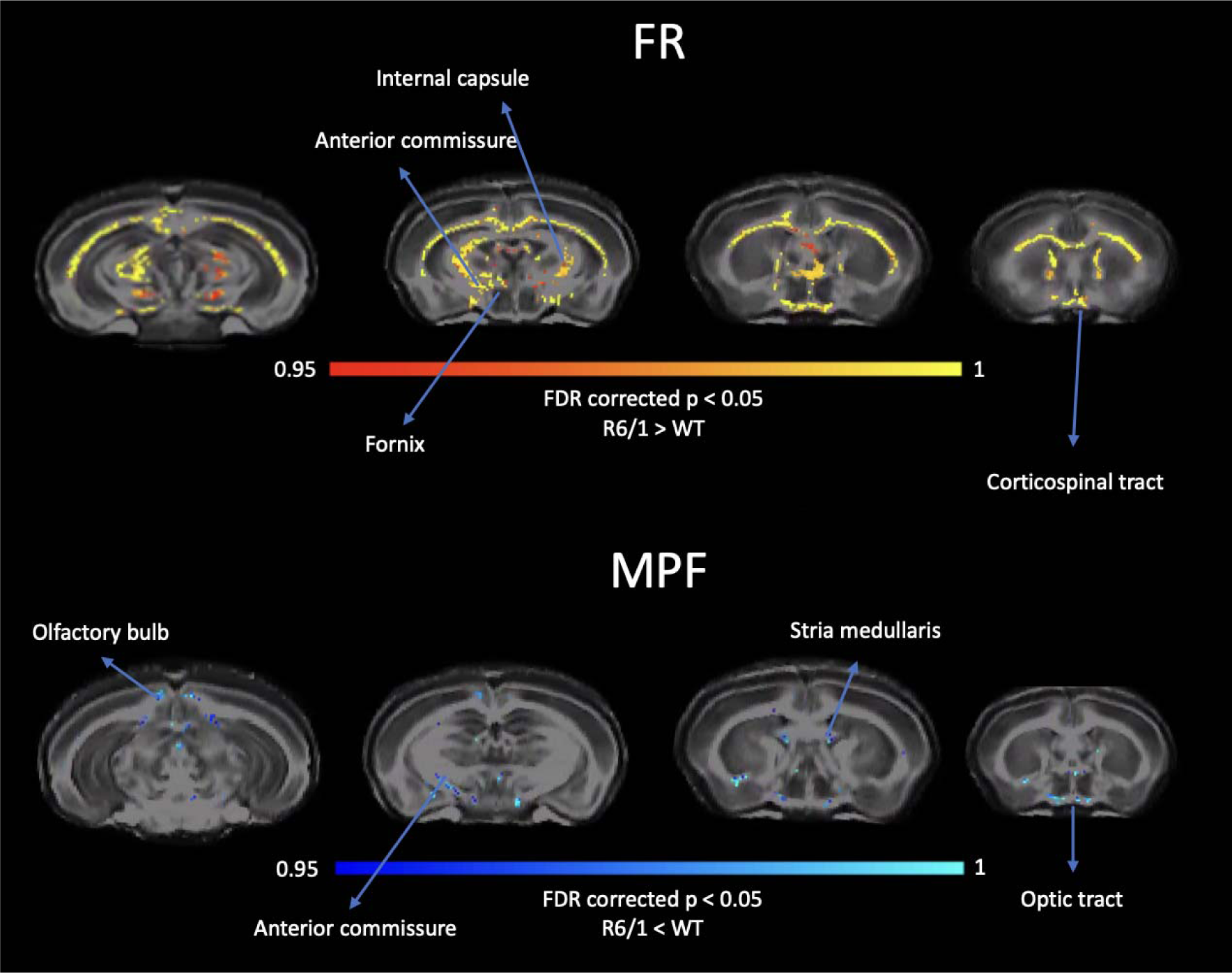
Tract-based spatial statistics (TBSS) analysis of white matter microstructure in R6/1 mice. White matter microstructural alterations were detected across the brain of R6/1 mice, revealing widespread increases in FR and some areas of decreased MPF. Abbreviations: FR: intra-axonal signal fraction. MPF: macromolecular proton fraction.

### 3.5. Increased axonal staining and decreased myelin staining in R6/1 mice

Representative images of MBP and NFL immunostaining in wildtype and R6/1 mice (Figure 6A-C). Across the medial-lateral axis, a higher percentage of CC area was immunostained for neurofilament light in R6/1 mice, as compared to wild-type mice, in the genu, body and splenium [Figure 6D-F; Splenium, min. F(1, 16)=5.00, p<0.05]. Conversely, a lower percentage area was stained for MBP in the R6/1 mice, as compared to wildtype mice, in the genu and body [Figure 6G-I; Genu, min. (F(1, 18)=6.03, p<0.05].

**Figure 6:**
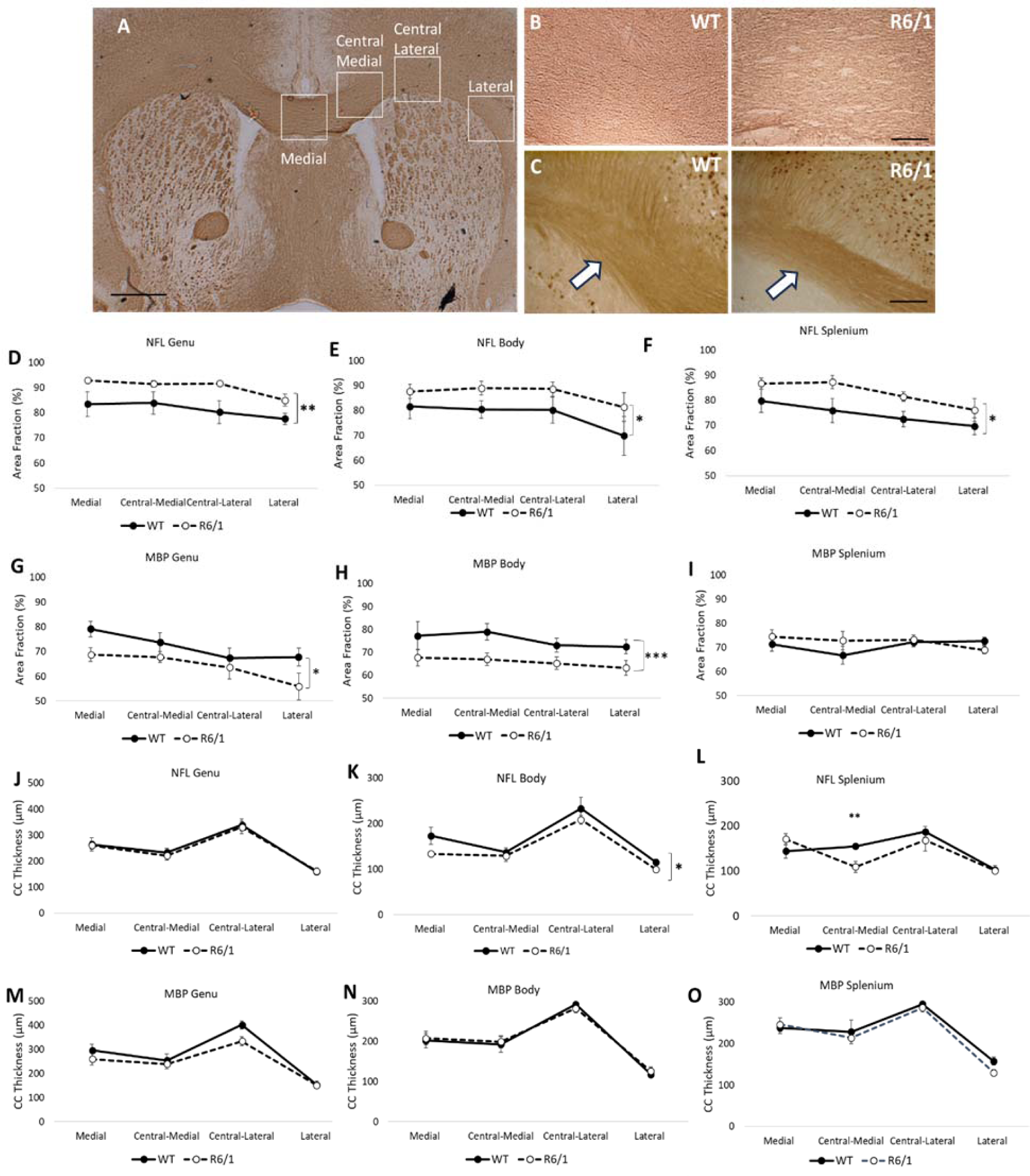
Immunohistochemical analysis of neurofilament light chain (NFL) and myelin basic protein (MBP) in WT and R6/1 mice. (A) Low magnification (x12.5) MBP immunostaining showing ROIs for analysis along the medial-lateral axis, with one medial ROI and three lateralised ROIs (Central-Medial, Central-Lateral, Lateral). Scale bar = 500µm. (B) Representative MBP immunostaining at high magnification (x100) in central-lateral region of the CC body for WT and R6/1 mice. Staining encompasses whole image. Scale bar = 200µm. (C) Representative NFL immunostaining at high magnification in the central-medial region of the CC body. White arrows indicate staining in CC. Scale bar = 200µm. Analysis of the percentage area stained (area fraction) for the NFL stain within the genu (D), body (E) and splenium (F). Analysis of the percentage area stained (area fraction) for the MBP stain within the genu (G), body (H) and splenium (I). Analysis of the CC thickness for the NFL stain within the genu (J), body (K) and splenium (L). Analysis of the CC thickness for the MBP stain within the genu (M), body (N) and splenium (O). Data are presented as mean ± S.E.M.

### 3.6. Thinner CC in R6/1 body based on NFL, but no difference in MBP

For the NFL immunostain, no differences in CC thickness were observed in the genu [Figure 6J; F(1,17)=1.63, n.s.], but the R6/1 mice had thinner CC measurements in the body across the medial-lateral axis [Figure 6K; F(1, 13)=6.66, p<0.05] and in the central-medial region of the splenium [Figure 6L; F(3, 48=3.47, p<0.05]. No differences in CC thickness were detected based on the MBP immunostain [Figure 6 M-O].

### 3.7. CC consists of thinner axons in R6/1 mice

Representative electron microscopy images from wildtype and R6/1 mice are shown in Figure 7A. A reduced g-ratio was calculated for R6/1 mice compared to wildtype [Figure 7C; t(4)=2.74, p=0.05]. Analysis of the raw data revealed a thinner mean axonal diameter [Figure 7D; t(4)=3.31, p<0.05] and this is reflected in the greater frequency of thinner diameter axons in R6/1 mice [Figure 7F]. No difference in myelin thickness was observed [Figure 7E; t(4)= −0.58, n.s.].

**Figure 7.**
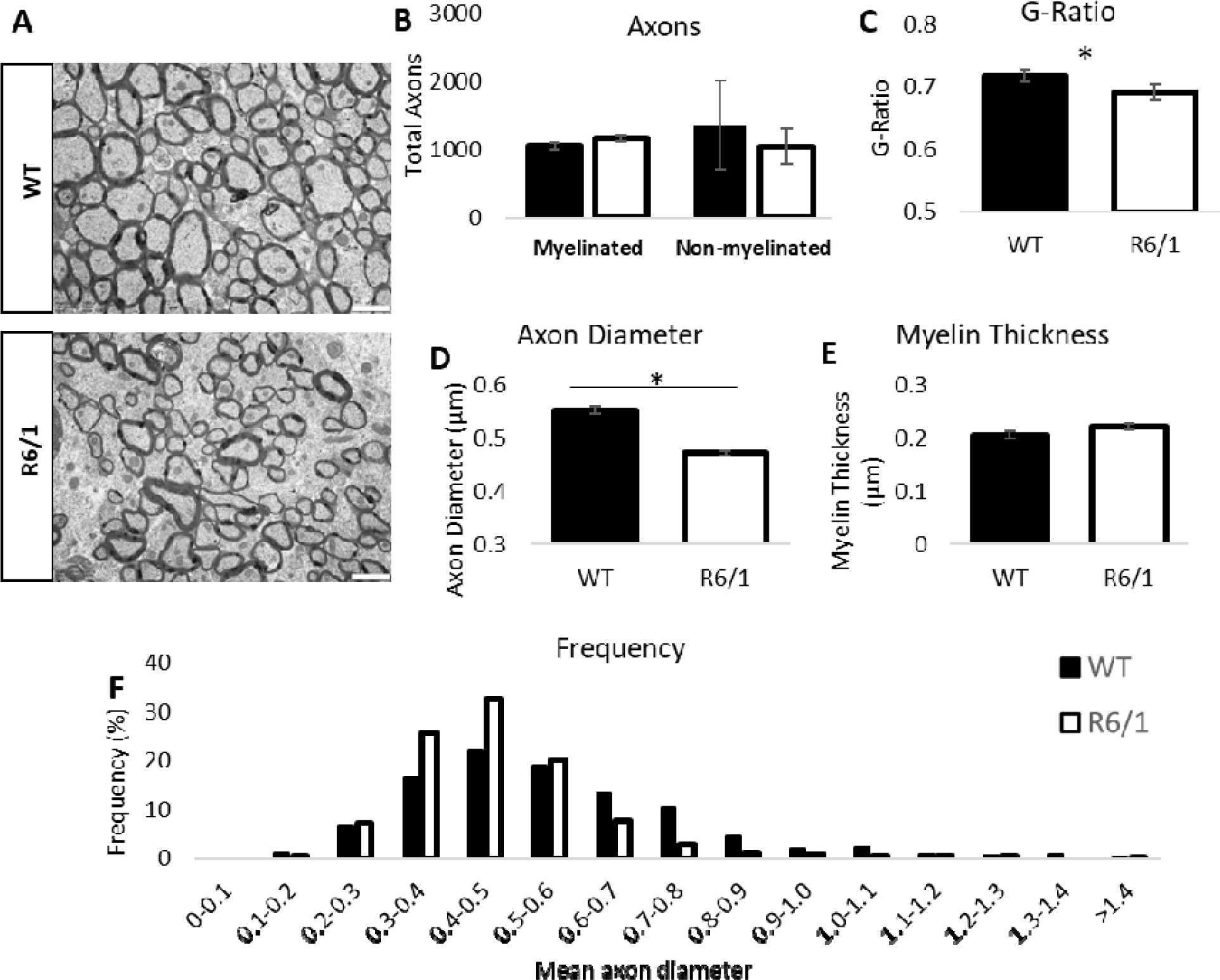
Analysis of axons and myelin via electron microscopy. (A) Representative TEM images from WT and R6/1 mice. White scale bars=1µm. (B) Total myelinated and non-myelinated axons in wildtype and R6/1 mice. (C) Mean g-ratio measurements for wildtype and R6/1 mice. (D) Mean axon diameter and (E) mean myelin thickness. (F) Percentage frequency of axons measuring 0.0- >1.4 µm in wildtype and R6/1 mice. N=3/group. Data are presented as mean ± S.E.M.

## 3 Discussion

We carried out a high-resolution *ex vivo* MRI assessment of WM alterations in the R6/1 mouse model of HD. We assessed 4 month-old mice to represent the early symptomatic stage of the disease ^40,42^. In order to explore the cellular causes underlying the imaging phenotype, histological and electron microscopy analyses were performed in a separate cohort of age- and sex-matched mice. Additionally, we investigated the functional correlates of cellular changes by assessing cognitive and motor function in a third age-matched cohort. We report motor and cognitive impairments, as well as novel findings of WM changes in this model of HD, which may inform future research in the human condition.

The motor and cognitive impairments detected in 4-month old R6/1 mice in this study are consistent with previous evidence ^40,42^. Interestingly, the cognitive deficits in response accuracy in the 5-CSRTT emerged early in training, even when the stimulus duration was long (10s) and the cognitive demands were low. This suggests that rather than an attentional deficit, R6/1 mice manifested a more global visuospatial impairment which disrupts their ability to code their responses in space. Although R6/1 mice were slower to respond, omitted more responses and completed fewer trials, they did not show any impulsivity. Additionally, they showed motor coordination, locomotor activity and dexterity impairments at 4 months old, but muscle strength was not affected. Given this disease-related functional impairment, we used high-resolution *ex vivo* imaging to explore the neurobiological differences between the genotypes.

MRI assessment of WM microstructural changes across the CC using a tractometry approach ^79,80^ revealed widespread increases in the intra-axonal signal fraction (FR) and decreased apparent myelin (MPF) in R6/1 mice. Interestingly, TBSS analysis uncovered widespread whole brain increases in FR, together with some regional decreases in MPF, suggesting that such alterations extend beyond the callosum. Importantly, the widespread FR increases detected in this study are consistent with findings from a recent MRI study on pre-symptomatic HD recently published by our group ^5^, which demonstrated that FR is increased in the cortico-spinal tract of premanifest HD patients. This points to the potential of FR as a cross-species MRI marker of premanifest axonal changes in HD.

It has to be mentioned that the CHARMED model ^39^ utilised in this study to model the diffusion signal and obtain FR maps represents axons as parallel cylinders and thus cannot recover the effect of axonal orientation dispersion due to bending and fanning of axon bundles widespread throughout the brain ^81^. More recent models such as the neurite orientation dispersion and density imaging (NODDI) model ^82^ support a description of WM beyond the most coherently-oriented structures and provide an estimate of the ‘spread’ of neurite orientations within a voxel with the orientation dispersion index (ODI).

Notably, a recent *ex vivo* MRI study by Gatto and colleagues ^42^ used NODDI to evaluate early microstructural changes in R6/1 mice, and reported increased ODI in the callosal genu at 11 weeks of age. The isotropic volume fraction (IsoVF), representing the signal coming from cerebrospinal fluid and other non-neurite tissue components, was also increased. On the other hand, the intracellular volume fraction (ICVF), representing the fraction of the MRI signal coming from the non-CSF compartment, was lower.

Based on the evidence provided by Gatto et al. ^42^ in combination with findings from the current study, we suggest that R6/1 mice present alterations in axonal morphology and structural organization, as reflected by more densely packed axons. Specifically, consistent with previous evidence from the R6/2 model ^83^, R6/1 mice likely suffer from a reduction in the overall density of neurites, which include axons, dendrites and other neuronal processes, as indicated by lower IsoVF and ODI ^42^. These neurites alterations in turn allow for a denser packing of the axons, leading to the higher FR detected in this study and in premanifest patients ^5^, as well as to the increased axon immunostaining detected across the callosum in this study.

Notably, we also show that R6/1 mice present axons with disproportionately small diameters, as demonstrated by a thinner mean axonal diameter, and a greater frequency of thinner diameter axons detected with electron microscopy.

Previous evidence has shown an inverse relationship between axon diameter and axon density. Specifically, larger diameter axons take up more space, due not only to larger size, but also from the space demands of neighbouring glial cells ^84–86^. Therefore, when there are fewer large diameter axons as shown here, or their morphology is less complex ^42,83^, the space requirements go down. As a consequence, the *relative* proportion of smaller diameter axons increases, leading to increases in the intra-axonal signal fraction and decreases in WM volume ^87^. Overall, this is consistent with our findings of thinner callosal splenium as detected with axonal immunostaining, as well as the increases in FR detected in the MRI sample.

It is possible that axons in this model develop normally but then shrink because of the disease process. However, it might also be that R6/1 mice present abnormalities in the postnatal development of axons and studies evaluating axon microstructure in R6/1 mice earlier in development will be useful in assessing such possibility. Consistent with a developmental effect of the mutation, evidence shows that neurotypical development is associated with an increase in thick axons relative to thinner ones, which in turn tend to be present in a higher proportion early in life ^88^.

In terms of myelin-related alterations in this model, histological analysis revealed that myelin staining was decreased in the genu and body, paralleling MPF decreases observed in the MRI sample. Interestingly, while MBP staining was decreased, electron microscopy did not detect significant alterations in the thickness of myelin sheaths in R6/1 mice.

Electron microscopy primarily focuses on the physical structure of myelin, such as the compactness of myelin layers, while MBP staining specifically targets the presence of MBP, which is a major component of myelin. This in turn suggests that, at least early in disease progression, R6/1 present a reduction in the expression or content of myelin-associated proteins, without significant alterations in the structure of myelin sheaths themselves. Such reductions might in turn be due to changes in gene expression, protein synthesis, or protein degradation.

Importantly, we detected increased WM volume in R6/1 mice with VBM. Interestingly, such an increase also concerned the splenium of the callosum, where NFL staining detected reduced thickness. Although these findings might seem counterintuitive, it is important to stress that VBM and NFL staining have different spatial resolutions and sensitivities. Specifically, while NFL staining provides fine-grained information about specific axonal changes, VBM captures broader structural changes that include factors other than axons. Therefore, an increase in volume as detected by VBM could be due to changes in the size or quantity of brain tissue components other than axons, such as glial cells, blood vessels, or inflammation-related oedema and swelling.

Neuroinflammation is one of the central mechanisms involved in HD neuronal death, and reactive inflammatory processes are one of the most studied at the early stages of HD ^89^. Accordingly, Gatto et al. ^42^ demonstrated the presence of neuroinflammatory processes from the early stage of the disease in R6/1 mice. Therefore, swelling of glial cells may have biased brain atrophy measurements ^90^ and may underlie the volume increases observed in this study. Inflammation-related swelling of glia may also explain the widespread increases FR observed in R6/1 mice, by reducing extracellular fluid and thus causing reduced diffusion in the extra-axonal space ^91,92^.

To summarise, we report differences in WM microstructure in the R6/1 mouse model of HD. By combining insight from MRI, histology and electron microscopy, as well as evidence from previous studies, we suggest that such alterations are likely driven by axonal changes. Specifically, we show that R6/1 mice present disruptions in axonal morphology and organization. Furthermore, we show that, at least early in disease progression, R6/1 present a reduction in the expression or content of myelin-associated proteins, without significant alterations in the structure of myelin sheaths themselves. Finally, our findings indicate that neuroinflammation-driven glial and axonal swelling might also affect this mouse model early in disease progression.

These results extend previous findings on animal models and HD patients, all of which demonstrate alterations in WM microstructure as a significant feature of HD ^4,5,17,18,21,23,32,33,93–95^. Crucially, we demonstrate cross-species convergence in FR increases, as we recently showed that alterations in this measure are also present in premanifest HD patients ^5^. This points to the potential of FR as a novel MRI biomarker of HD-associated changes in WM microstructure.

Biomarkers constitute an important tool of translational science in medicine, thus the development of biomarkers that are useful and accessible in both animal models and humans affords improved translation of animal findings into humans and, likewise, the back-translation of imaging methods from humans to animal models ^96^. Therefore, while inherent differences between species remain, the present findings may represent an important preliminary step in the establishment of FR as a novel imaging biomarker of WM pathology in HD.

### Methodological considerations

The MRI data were obtained *ex vivo* from fixed tissue. As several factors, such as tissue fixation, sample temperature, and the acquisition scheme all have an effect on MRI measures, considerable care should be taken when extrapolating the present *ex vivo* findings to *in vivo* values ^97^. However, the observed group differences should be valid, since any bias in terms of methodology should have the same effect of the two groups compared. Furthermore, in the present study, several steps were taken with regards to the perfusion and tissue preparation protocols to obtain optimal tissue quality. Such procedures were based on previous literature ^45^, and consisted of the utilisation of liquid, rather than powder, perfusates, as well as delivery of perfusate at a low flow rate, in order to avoid blockage of the capillary beds and vessels’ rupture. Additionally, the storage of tissue in a sodium azide solution enabled improved tissue conservation.

Another important thing to consider is that this study assessed three distinct cohorts of mice. Therefore, the nature of our study design limits our ability to establish causal relationships between variables. Nevertheless, our study provides an important stepping stone for future research assessing axonal alterations as a pathophysiological feature of HD.

## Acknowledgements and funding source

The present research was funded by a Wellcome Trust PhD studentship to CC (ref: 204005/Z/16/Z). MJL and AER were funded by an MRC Programme grant (MR/T033428/1). MJL was funded by an MRC-AMED grant (MR/V00560X/1) and an NC3Rs grant (NC/X001016/1), as well as a Parkinson’s UK Fellowship (F-1502). DKJ was supported by a New Investigator Award (to DKJ) from the Wellcome Trust (ref: 096646/Z/11/Z) and a Strategic Award from the Wellcome Trust (ref: 104943/Z/14/Z). VD’s laboratory is supported by the UK Dementia Research Institute (DRI-TAP2022FA2 and UK DRI-3006) which receives its funding from UK DRI Ltd, funded by the UK Medical Research Council, Alzheimer’s Society and Alzheimer’s Research UK. VD is further supported by a professorship from the Academy of Medical Sciences (AMSPR1\1014).

## Data sharing and availability

The data analysed during the current study and the respective analysis scripts are available from the corresponding author on reasonable request.

